# Dimorphic development in *Streblospio benedicti:* genetic analysis of morphological differences between larval types

**DOI:** 10.1101/008730

**Authors:** Christina Zakas, Matthew V. Rockman

## Abstract

The marine polychaete *Streblospio benedicti* exhibits two distinct larval types, making it a model for the study of developmental evolution. Females produce either large eggs or small ones, which develop into distinct lecithotrophic or planktotrophic larvae with concomitant morphological and life-history differences. Here, we investigate the inheritance of key morphological traits that distinguish the larval types. We use genetic crosses to establish the influence of maternal and zygotic differences on larval phenotypes. We find a large maternal effect on larval size and the number of larval chaetae, while the number and length of these chaetae are also strongly influenced by zygotic genotype. Interestingly, the distribution of larval phenotypes produced by these crosses suggests traits intermediate to the two parental types should not be uncommon. Yet, despite gene flow between the types in natural populations, such intermediates are rarely found in nature, suggesting that selection may be maintaining distinct larval modes.

## Introduction

The evolution of development couples molecular and cytological processes with ecological and population-genetic phenomena such as dispersal and life-history strategies. Benthic marine animals are particularly well suited to studies of developmental evolution in an ecological context, because while lineages exhibit a broad diversity of adult forms, their pelagic larvae fit loosely within one of two suites of correlated life-history characteristics (Thorson, 1950): *planktotrophic* larvae are small, obligately feeding larvae that spend days to months developing in the plankton before reaching metamorphic competence; *lecithotrophic* larvae generally lack complex feeding structures and are able to metamorphose and settle to the benthos soon after their release, provisioned only by maternal yolk (Strathmann, 1985).

How and why this developmental dichotomy occurs has been a subject of longstanding debate and inquiry (Marshall and Morgan, 2011; McEdward, 2000; Moran and McAlister, 2009; Pechenik, 1999; Vance, 1973). While divergent developmental modes are common across pairs of sister species (Strathmann, 1978; Wray and Raff, 1991), most species exhibit only one mode, precluding genetic analysis of differences between offspring morphs and confounding larval mode with all other interspecific differences. Little is known about how, at the population level, suites of life history traits related to larval mode vary and evolve.

The phenomenon of poecilogony, whereby a single species produces distinct types of embryos and thus two distinct life-history strategies, offers a route toward answering these questions (Knott and McHugh, 2012). Poecilogony provides a microevolutionary model for the pervasive macroevolutionary pattern of larval life-history evolution, enabling forward genetic dissection of developmental and life-history traits and their genetic correlations. Here we use the polychaete *Streblospio benedicti*, the best characterized case of poecilogony from a genetic perspective, to explore the genetic basis for several characters that distinguish planktotrophic and lecithotrophic larvae.

*S. benedicti* is a common spionid polychaete that inhabits muddy estuarine sediments. Adults live in the sediment and dispersal is primarily, if not exclusively, achieved in the larval phase. Females produce either clutches of tens of lecithotrophic embryos, each ∼200 µm diameter, or clutches of hundreds of planktotrophic embryos of, each ∼100 µm diameter (Bridges, 1993; Levin, 1984; Levin and Bridges, 1994; McCain, 2008). Classical quantitative genetic studies indicate that poecilogony is a heritable polymorphism in *S. benedicti,* and that larval mode is unaffected by large differences in temperature, day length, and food availability (Levin, 1986; Levin and Bridges, 1994; Levin and Creed, 1986; Levin *et al*., 1991). In particular, Levin *et al*. (1991) used a reciprocal mating design between adults of the two larval modes and found a large additive contribution for many reproductive and larval traits associated with life history mode.

Females of both offspring types can co-occur in small spatial areas, although generally populations tend to consist of only one offspring type (Zakas and Wares, 2012). Both mitochondrial and nuclear loci indicate recent gene flow between the two morphs (Rockman, 2012; Schulze *et al*., 2000; Zakas and Wares, 2012). The rare occurrence of contrasting suites of heritable larval traits within a single species provides an opportunity to determine the genetic components of phenotypic variance associated with life history.

Here we use experimental crosses to disentangle the roles of zygotic genetic effects and maternal effects. We use a mating design with replicate families of reciprocal crosses to identify the genetic components of phenotypic variation associated with the two larval types. We extend our crosses to F_2_ and four classes of backcross, and we describe, for the first time, larval morphological variation within genetically segregating families in *S. benedicti*.

We focus our study on a few key morphological differences between the two larval types. First, alternative types of *S. benedicti* larvae differ dramatically in size as a result of their difference in egg volume. We ask whether the anterior-posterior axis length of a larva depends only on the oocyte cytoplasm or whether it is influenced as well by the larval genome. Second, we focus on a striking morphological difference between the two types: the larval chaetae. Chaetae are epidermal structures made of β-chitin in a protein matrix, and are ancestral characters of annelids and name-defining for polychaetes (Hausen, 2005). Spionid larvae typically have many long larval chaetae protruding from two chaetal sacs on either side of the first segment. Though lecithotrophic larvae develop chaetal sacs, the larval chaetae themselves are absent (Gibson *et al*., 2010, and Fig. 1). Planktotrophic larvae spend a long period developing in the plankton where predation mortality is large and chaetae can confer an advantage by deterring predation (Pennington and Chia, 1984; Pernet *et al*., 2002; Vaughn and Allen, 2010). In lecithotrophic larvae, these chaetae may be unnecessary or disadvantageous.

**Figure 1.**
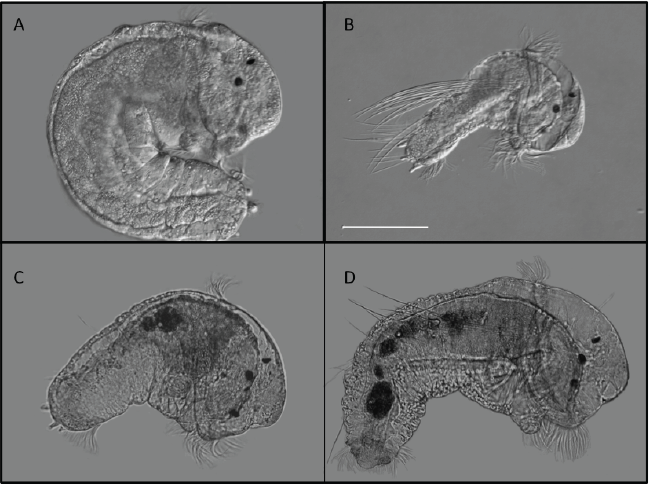
*S. benedicti* larvae upon release from the maternal brood pouches, all shown at the same magnification. **A-B,** DIC images of parental (wild-type) larvae. **A,** a Long Beach lecithotrophic larva (cross type LxL). **B**, a Bayonne planktotrophic larva (cross type BxB). **C-D,** brightfield views of representative F_2_ larvae. The larva in **C** has no chaetae originating from the chaetal sac, similar to the LxL larva in **A**. The white scale bar is 100 µm.

## Methods

### Crosses

Individuals derive from two populations. Lecithotrophic animals from Long Beach, CA, and planktotrophic animals from Bayonne, NJ, have been maintained under common conditions in the lab. The Long Beach larvae are lecithotrophic in the sense that they need not feed prior to settlement. However, they are facultatively planktotrophic, eating when fed (Pernet and McArthur, 2006; and personal observation). Both Bayonne and Long Beach have been sampled multiple times in previous years and the offspring mode recovered is consistent (Rockman, 2012; Zakas and Wares, 2012). Larvae from wild-collected adults of these populations were collected and planktotrophic larvae were fed an excess of *Dunaliella salina* algae until settlement. Lecithotrophic offspring were collected and transferred immediately to new sediment. Juveniles and adults were kept individually in wells of 6-well cell-culture plates with artificial seawater (Instant Ocean, 28 ppt in milliQ water) and ∼1ml defaunated mud slurry, which completely covers the bottom of the well. The mud averages 0.5 g dry weight per ml of slurry. The mud, collected from a tidal mudflat in Newark Bay, New Jersey, was sieved through 250µm mesh, frozen at −80°C, heated to 55°C, and then returned to room temperature for use. Water and mud were changed bi-weekly. Virgin females were placed with reproductive males (determined by the presence of spermatophores in the well) and checked daily for broods. Egg size was measured for a representative female of each type by removing oocytes or early embryos from brood pouches and imaging them on a slide at 20x magnification as described below for the larvae. Crosses were performed in both directions to assess maternal effects. Animals from a single F_1_ family were used as parents in the backcross and F_2_ families to minimize genetic variation. Each class of cross is represented by larvae from one to five full-sib families as listed in Table 1.

**Table 1.**
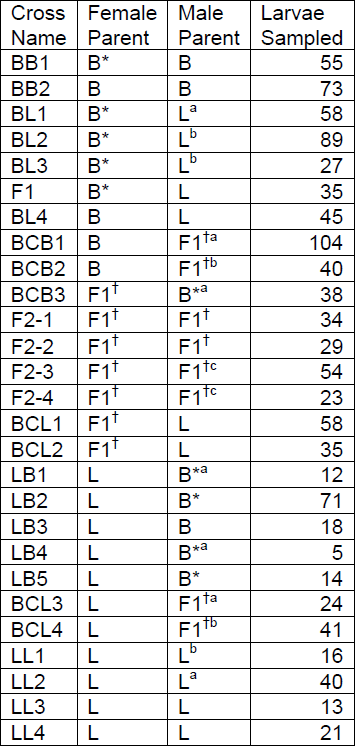
Experimental Crosses B = Bayonne, L = Long Beach. The asterisks indicate Bayonne individuals that are full siblings, and the crosses indicate F1 individuals that are full siblings. Superscripts a, b, and c indicate that the same male was used in multiple crosses.

## Measurements

Larvae were collected and measured at release from maternal brood pouch, prior to larval feeding, and imaged on a Zeiss Axio Imager A2 microscope with achromatic 10x and 20x dry objectives. Because larvae were measured prior to feeding, differences in phenotypes are due strictly to maternal and zygotic developmental factors. Pictures were taken of live larvae, which were slightly compressed under a coverslip on a depression slide coated with 400µl of 0.2% agarose in artificial seawater, which limited their movement and allowed for image capture (Fig. 1). Some larvae were fixed prior to measurement. These were relaxed in 7% MgCl_2_ and transferred to 4% formaldehyde in seawater overnight, followed by transfer to PBS. Comparative measurements between fixed and live larvae showed fixation did not have an effect on larval size or chaetal measurements. Measurements of larval gut length, chaetal length, and number of chaetae per chaetal sac were recorded in ImageJ (Schneider *et al*., 2012). To account for possible variation due to focal plane, three pictures of each larva were taken in different focal planes so that replicate measurements of each trait could be averaged.

Gut length was measured by tracing the dorsal edge of the gut in ImageJ. We averaged three replicate measurements (focal planes) for each larva, and we log-transformed these averages to generate a normally-distributed variable. The gut length dataset includes observations for 1,041 larvae. The number of larval chaetae was determined by counting chaetae on one side of each of 1,070 larvae. The length of the longest observed chaeta was studied in a dataset including only the 674 larvae that have chaetae.

### Fitting Models of Inheritance and Estimating Maternal and Zygotic Effects

With data on nine distinct classes of cross between lecithotrophic Long Beach animals (L) and planktotrophic Bayonne animals (B), we can separately estimate the effects of maternal and zygotic differences between the two populations on each phenotype. Our approach to this line-cross dataset follows the line-cross mean analysis laid out by Lynch (1991) and references therein, and described in Lynch and Walsh (1998, chapter 9), with modifications to incorporate maternal effects. Briefly, we model the mean larval phenotype of each class of cross as a function of the fraction of its genome that derives from the Bayonne population, the fraction of the animals in the line-cross that are expected to be heterozygous for alleles that differ between B and L, and the type of egg that the larva develops from (Fig. 2).

**Figure 2.**
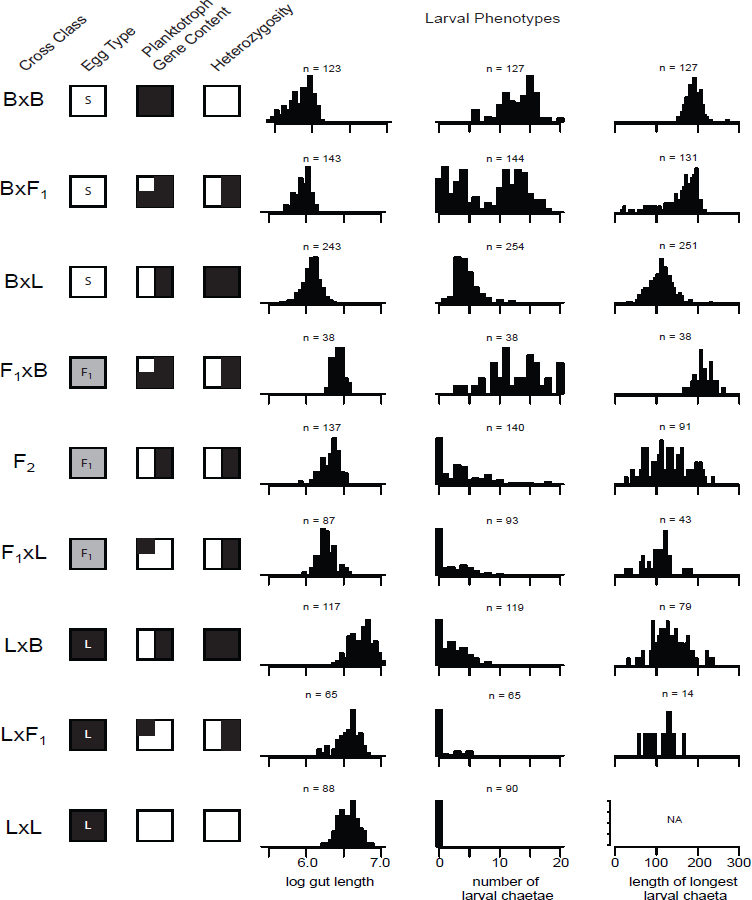
Larval phenotype distributions as a function of maternal and zygotic constitution. Each row represents a class of cross. The crosses are grouped into three sets by *Egg Type*: small eggs, F_1_ eggs, and large eggs. Within each egg type category, crosses are ordered by *Bayonne Gene Content*. The cross classes also vary in the proportion of individuals that are heterozygous for alleles from the two populations, as shown in the *Heterozygosity* column. The number (n) of individual larvae measured for each phenotype is reported above each histogram.

To isolate line-cross means from family-level effects (e.g., shared environment), we used mixed-effect linear models, incorporating family as a random effect. Family includes maternal effects that are unique to individual females. We estimate the composite additive effect of zygotic gene action, *α_Z,_* by regressing phenotype on each family’s Bayonne gene content (the source index, *θ_S_*), using the F_2_ population as a reference point. In this formulation, *θ_S_* is 0 in the F_2_ and other populations that carry equal proportions of Bayonne and Long Beach alleles, −1 in pure Long Beach animals, 1 in pure Bayonne animals, and −0.5 and 0.5 in each backcross. We extend this regression approach to estimate dominance effects, *δ_Z_*. The expected fraction of heterozygotes in each population, relative to the F_2_ population, is the variable *θ_H_* (the hybridity index). In this formulation, F_2_ and backcross populations, which are expected to be ½ heterozygous, have a hybridity index of 0, while F_1_s (completely heterozygous) have an index of 1 and the parental populations have an index of −1.

To estimate maternal effects that are due to egg type, we modeled an additive maternal effect, *α_M_*. Larvae derived from large eggs carry an *α_M_* coefficient (which we term *θ_MA_*) of 1, F_1_ eggs 0, and small eggs −1. To accommodate nonadditivity (i.e., F_1_ eggs conferring maternal effects that are not perfectly intermediate to large and small eggs), we also considered a dominant maternal effect, *δ_M_*. Its coefficient (*θ_MD_*) is 1 in F_1_-derived eggs and 0 in large and small eggs. Coefficients for the nine cross class line means are shown in Table 2 and schematized in Fig. 2.

We examined nested models incorporating the random effect of family only (null) and successively adding the additive and dominant maternal and zygotic effects, *α_M_*, *α_Z_*, *δ_Z_*, and *δ_M_* in the order of their anticipated effect size. The complete model for a measured phenotype of *y* of larva *j* in family *i* is

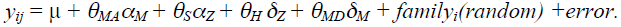

**Table 2.**
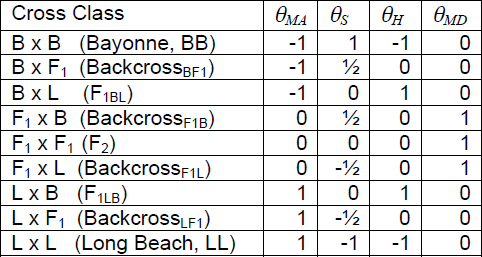
Coefficients for analysis of line-cross means In each cross class, the first named line is the mother.

All analyses were performed in R (R Core Development Team, 2011), version 2.14.1. Models for the chaetal length and log gut length phenotypes were fit by REML using the R function *lmer* (Bates *et al*., 2011). The chaetal number phenotype includes a large number of observations of zero chaetae, and the data are not conducive to analysis with Gaussian assumptions about error. We instead fit a Poisson generalized linear mixed-effect model using the *glmer* function, proceeding otherwise as above. Model fit was tested by comparing log likelihoods with a chi-square test. We also report alternative model-fit criteria (AIC and BIC), which support identical conclusions.

The maternal effects are themselves heritable, as both Bayonne planktotrophs and Long Beach lecithotrophs breed true in the common environment in the lab (and as documented by Levin *et al*., 1991). However, our crosses do not allow segregation of egg-size or other maternal effects, as we observed larvae only through F_2_ and backcross generations. As a result, maternal effects are correlated with zygotic effects; across the three egg size classes, larvae derived from planktotrophic eggs have more Bayonne genes than larvae derived from lecithotrophic eggs (see Fig. 2). Consequently, we can only compare zygotic and maternal effects when both terms are simultaneously included in our models. Reported effect estimates therefore derive from the full model, even in cases where individual terms did not achieve formal significance.

We estimated the effective number of segregating factors affecting larval chaeta length using Lande’s (1981) modifications of the Castle-Wright estimator (Wright, 1968), adapted to the peculiarities of the phenotype as described in the results.

## Results

Egg size, estimated as the average of two orthogonal measurements of the diameter, falls into three discrete classes. The eggs of Bayonne planktotrophs averaged 102 µm, with a standard deviation of 7.5 µm (n = 8), those of Long Beach lecithotrophs averaged 205.1 ± 4.9 µm, (n=8) and the F_1_ eggs averaged 137.1 ± 5.4 µm (n=30). The planktotrophic and lecithotrophic eggs thus differ in volume by a factor of 8, and the F_1_ eggs are closer in volume to those of the planktotrophs than to those of the lecithotrophs.

Inheritance of larval body length at release, measured as larval gut length, is almost completely governed by maternal effects (Fig. 2). The nine classes of line cross are ordered in the figure so that the first three derive from small eggs, the next three from F_1_ eggs, and the last three from large eggs. Within each of these groups of three, the crosses are ordered from most Bayonne genome content to least. Statistical analysis (Table 3) confirms the graphical impression from Fig. 2, that the three sets of three differ substantially from one another, with smaller differences within each set. The best-fitting model incorporates additive maternal effects and additive and dominant zygotic effects. However, additive zygotic effects are not significant on their own and the zygotic effects are small in magnitude. As suggested by Fig. 2, the direction of the zygotic effects is inconsistent between the egg-size classes, suggesting that cytoplasmic factors may modulate zygotic effects. An alternative model parameterization that includes *α_M_*, *α_Z_*, and an explicit maternal-by-zygotic interaction term (*α_M_* x *α_Z_*) fits almost as well as the model with *α_M_*, *α_Z_*, and *δ_Z_* (Table 3). We can conclude that zygotic gene action influences larval gut length, but that little of that influence is additive, at least in composite across the genome.

**Table 3.**
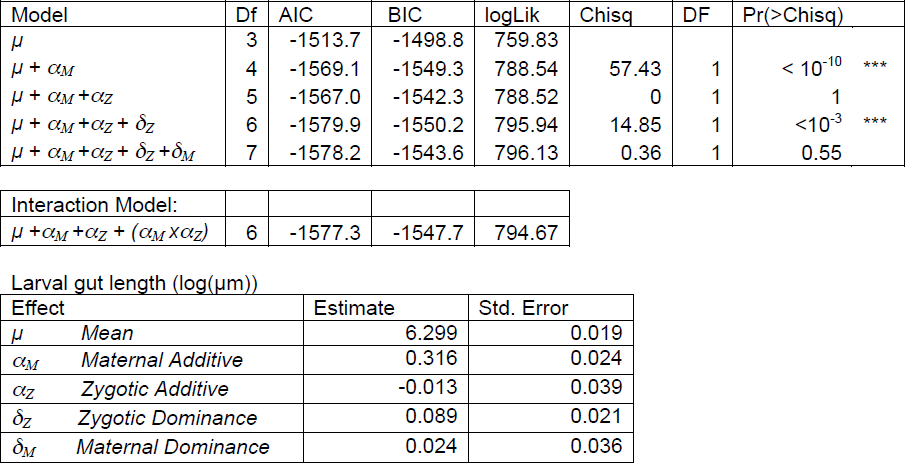
Larval gut length model fit and effect estimates

| Model | Df | AIC | BIC | logLik | Chisq | DF | Pr(>Chisq) |
| --- | --- | --- | --- | --- | --- | --- | --- |
| $\mu$ | 3 | -1513.7 | -1498.8 | 759.83 | | | |
| $\mu + \alpha_M$ | 4 | -1569.1 | -1549.3 | 788.54 | 57.43 | 1 | $< 10^{-10}$ *** |
| $\mu + \alpha_M + \alpha_Z$ | 5 | -1567.0 | -1542.3 | 788.52 | 0 | 1 | 1 |
| $\mu + \alpha_M + \alpha_Z + \delta_Z$ | 6 | -1579.9 | -1550.2 | 795.94 | 14.85 | 1 | $< 10^{-3}$ *** |
| $\mu + \alpha_M + \alpha_Z + \delta_Z + \delta_M$ | 7 | -1578.2 | -1543.6 | 796.13 | 0.36 | 1 | 0.55 |

|  |  |  |  |  |
| --- | --- | --- | --- | --- |
| Interaction Model: |  |  |  |  |
| $\mu + \alpha_M + \alpha_Z + (\alpha_M \times \alpha_Z)$ | 6 | -1577.3 | -1547.7 | 794.67 |

Larval gut length (log( μm))
| Effect | Estimate | Std. Error |
| --- | --- | --- |
| $\mu$ Mean | 6.299 | 0.019 |
| $\alpha_M$ Maternal Additive | 0.316 | 0.024 |
| $\alpha_Z$ Zygotic Additive | -0.013 | 0.039 |
| $\delta_Z$ Zygotic Dominance | 0.089 | 0.021 |
| $\delta_M$ Maternal Dominance | 0.024 | 0.036 |

Inheritance of larval chaetae number is markedly different (Fig 2). Zygotic effects are pronounced. Within each of the egg-size classes, increasing proportions of Bayonne alleles increase chaetae number. However, maternal effects are also pronounced. This is clear from contrasts between reciprocal crosses that differ only in egg size (F_1BL_ vs F_1LB_, Backcross_BF1_ vs Backcross_F1B_, Backcross_LF1_ vs Backcross_F1L_). Segregation variance is visible in these data, as more variation in chaetae number is present in the backcross and F_2_ classes due to genetic variation, as discussed below. Variation in penetrance is also evident in chaetae number: within F_1LB_ families all larvae are identical in their Bayonne genetic content, but only 30% are lacking chaetae. Our full model, incorporating additive and dominance maternal and zygotic effects, is favored by the data, with additive zygotic effects the largest (Table 4).

**Table 4.** Larval chaetae number model fit and effect estimates

| Model | Df | AIC | BIC | logLik | Chisq | DF | Pr(>Chisq) |
| --- | --- | --- | --- | --- | --- | --- | --- |
| $\mu$ | 2 | 2435.6 | 2445.5 | -1215.8 | | | |
| $\mu + \alpha_M$ | 3 | 2417.8 | 2432.7 | -1205.9 | 19.76 | 1 | $<10^{-5}$ *** |
| $\mu + \alpha_M + \alpha_Z$ | 4 | 2390.4 | 2410.3 | -1191.2 | 29.39 | 1 | $<10^{-7}$ *** |
| $\mu + \alpha_M + \alpha_Z + \delta_Z$ | 5 | 2384.1 | 2409.0 | -1187.1 | 8.30 | 1 | 0.004 ** |
| $\mu + \alpha_M + \alpha_Z + \delta_Z + \delta_M$ | 6 | 2365.2 | 2395.1 | -1176.6 | 20.90 | 1 | $<10^{-5}$ *** |

Number of chaetae (Poisson mean)
| Effect | Estimate | Std. Error |
| --- | --- | --- |
| $\mu$ *Mean* | 0.509 | 0.103 |
| $\alpha_M$ *Maternal Additive* | -0.464 | 0.099 |
| $\alpha_Z$ *Zygotic Additive* | 2.126 | 0.203 |
| $\delta_Z$ *Zygotic Dominance* | 0.538 | 0.117 |
| $\delta_M$ *Maternal Dominance* | 0.856 | 0.161 |

We modeled the length of the longest measured chaeta for each larva, excluding zero-chaetae animals from consideration. We found no evidence of maternal effects, but there is strong support for additive zygotic effects (Table 5; *p* < 10^−5^). Thus our data suggest that chaetal length is inherited zygotically, but that the penetrance of chaetal growth (i.e., the presence or absence of chaetae) depends on maternal effects as well.

**Table 5.** Longest larval chaeta model fit and effect estimates

| Model | Df | AIC | BIC | logLik | Chisq | DF | Pr(>Chisq) |
| --- | --- | --- | --- | --- | --- | --- | --- |
| $\mu$ | 3 | 7624.6 | 7638.6 | -3809.3 | | | |
| $\mu + \alpha_M$ | 4 | 7625.1 | 7643.7 | -3808.5 | 1.56 | 1 | 0.21 |
| $\mu + \alpha_M + \alpha_Z$ | 5 | 7605.4 | 7628.6 | -3797.7 | 21.68 | 1 | $<10^{-5}$ *** |
| $\mu + \alpha_M + \alpha_Z + \delta_Z$ | 6 | 7606.3 | 7634.2 | -3797.2 | 1.07 | 1 | 0.30 |
| $\mu + \alpha_M + \alpha_Z + \delta_Z + \delta_M$ | 7 | 7607.4 | 7639.9 | -3796.7 | 0.94 | 1 | 0.33 |

Length of longest chaeta ( $\mu\text{m}$ )
| Effect | Estimate | Std. Error |
| --- | --- | --- |
| $\mu$ *Mean* | 124.879 | 8.698 |
| $\alpha_M$ *Maternal Additive* | 7.916 | 6.691 |
| $\alpha_Z$ *Zygotic Additive* | 77.377 | 16.996 |
| $\delta_Z$ *Zygotic Dominance* | -3.120 | 10.013 |
| $\delta_M$ *Maternal Dominance* | 12.175 | 12.546 |

Moreover, conditional on chaetal presence, number and length are correlated. In Fig. 3A, we show the correlation between the residual variation in chaetal number and the residual variation in chaetal length, after the effects of *α_M_*, *α_Z_*, *δ_Z_*, *δ_M_,* and family have been accounted for. This correlation (*r* = 0.49, p < 10^−15^) may be due to the segregation of genetic factors within families or to common microenvironmental effects influencing both traits in individual larvae. Tentative evidence for genetic correlation comes from the generally higher correlation among the residuals of F_2_s and Backcross populations, which are genetically heterogeneous with respect to alleles that differ between Bayonne and Long Beach, than among the residuals of the other populations, which are genetically homogeneous (Fig. 3B).

**Figure 3.**
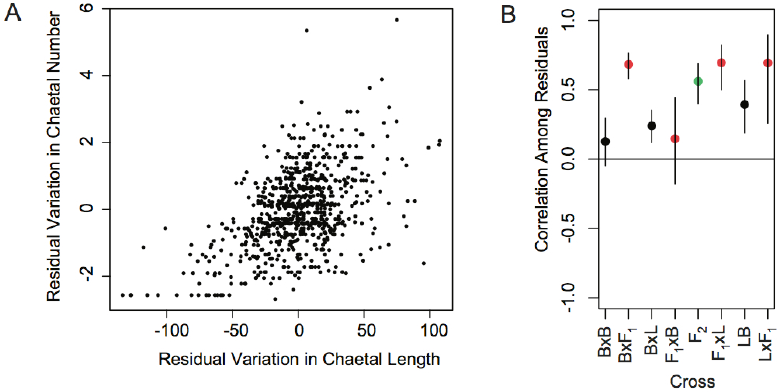
The length and number of larval chaetae are correlated within families. **A**, the distribution of residual chaetae length and number for all larvae. **B,** mean and confidence intervals of the correlation among residuals within each class of cross. The correlation coefficients are generally higher in populations in which two genotypes are segregating at a locus (backcrosses, red) or three genotypes (F_2_s, green) than for populations that are not segregating (parentals and F_1_s, black). The LxL cross is not shown because it lacks larval chaetae.

Given the additive zygotic nature of the inheritance of chaetal length, we can employ classical biometrical techniques to crudely estimate the underlying genetic architecture. Although the absence of chaetae from one of the parental classes complicates the analyses, Lande (1981) showed that segregation variance, σ*_s_*^2^, can be estimated from the phenotypic variances of F_2_ and reciprocal backcross populations:

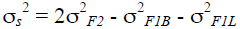

Note that these three classes of larvae occur on the same maternal-effect background (F_1_ oocytes). We estimated variances for these three line-cross classes using the residuals of the linear mixed-effect model. The effective number of factors (independently segregating genes of equivalent effect size) accounting for this amount of segregation variance (2,679 µm^2^) can be estimated by

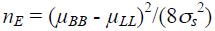

where BB and LL are the parental (Bayonne and Long Beach) crosses. Here, µ*_LL_* is unknown, because the lecithotrophic larvae lack chaetae. We can use an alternative estimator for the parental phenotypic difference: twice the additive effect of Bayonne alleles, as estimated from our linear model. That estimate gives a predicted Bayonne chaetal length of 202.3 µm and a predicted Long Beach chaetal length of 47.5 µm (i.e., if Long Beach larvae had chaetae; though theoretical, this is the chaetal length the model predicts for a Long Beach genome in a Bayonne egg). We conclude that, conditional on the presence of chaetae, chaetal length is influenced by 1.12 additive genetic factors.

## Discussion

The three larval developmental traits we measured differ in their underlying genetic architecture. Larval gut length is almost entirely maternally determined with a small, non-additive zygotic contribution. The presence of larval chaetae and their number involve substantial effects of both egg type and larval genome. Remarkably, the length of the chaetae — in larvae that have chaetae — appears to depend exclusively on additive zygotic effects. The genetic contribution to chaetae number and length is also correlated within families; larvae with long chaetae tend to have many (>10) while larvae with short chaetae only have a few (1-5).

We find that larvae originating from the same egg class are generally of the same size at release, as assessed by body length measured along the gut. F_1_ mothers produce larvae that are intermediate in size to the two parental lines (Fig. 2). Although lecithotrophic gut development is delayed by about a day relative to planktotrophs (Pernet and McHugh, 2010), the gut is fully formed before release in both larval morphs. Small differences in gut length within egg-size classes relate to features of the zygotic genome, but the direction of this zygotic effect is not consistent across egg size classes. This may be due to greater variability in the larval stage, and thus size, at which lecithotropic females released their larvae (results not shown). As larvae age, they increase the number of setigers and begin to develop adult features such as branchiae and palps. We observed that the same lecithotrophic mother could release subsequent broods at earlier or later stages, although larvae within a release were at the same developmental stage. This difference in developmental stage at release could be due to microenvironmental differences between clutches (such as longer days or temperature variation), random chance, or a response to physical perturbance. While lecithotrophic mothers showed variation in larval sizes across clutches, larvae from F_1_ mothers tended to have variability in larval size within a clutch, which was especially noticeable in the F_2_s (Fig. 1 C-D). Planktotrophic mothers, on the other hand, released larvae at the same stage consistently.

We observe that a large egg does not necessarily result in a “phenotypically lecithotrophic” offspring lacking chaetae. While the size of a larva is almost entirely maternally specified, there is variation in the penetrance of chaetal development. Development of the larval chaetae begins a few days after fertilization, consistent with an important role for zygotic gene activity, and the chaetae are fully formed by the time the larvae are released. While chaetal sacs occur in all larvae, the presence and number of chaetoblasts, the cells responsible for generating chaetae, are not known for each morph. Approximately 70% of F_1LB_ larvae develop some chaetae, though these larvae all carry one copy each of the Long Beach and Bayonne genomes. Further, ∼23% of Backcross_LF1_ larvae, which are homozygous for Long Beach alleles across 50% of the genome and heterozygous at the other 50%, also develop chaetae. Additionally, our estimate of the additive affect of Bayonne alleles on chaetal length suggests that a larva with a pure Long Beach genome in a small egg would produce moderate-length chaetae. This indicates that a switch in both maternal specification and zygotic gene action has to occur to produce a phenotypically lecithotrophic larva from a planktotrophic background. This pattern is particularly interesting in the context of life-history evolution from small to large egged larvae that has been described in most marine invertebrate groups. It demonstrates that simply increasing maternal allocation to egg size is not sufficient to produce a phenotypically lecithotrophic larvae.

Previous work in *S. benedicti* showed that many of the traits associated with life history strategy are highly heritable. Levin et al. (1991) estimated heritability from crosses between populations of planktotrophic and lecithotrophic parents within Bogue Sound, NC. While our phenotypes also show a strong genetic component, our results are not directly comparable to theirs as the only overlapping measurement is presence of larval chaetae. However, Levin et al. (1991) measured a different phenotype: proportion of families within a cross that had chaetae present, whereas we measured variation in chaetae number within a family. They found that the presence of chaetae is highly heritable (93.9%), but they did not recover a significant maternal effect for chaetal presence.

*S. benedicti* evolved poecilogony from an ancestor with a single life-history strategy, most likely planktotrophy (Mahon *et al*., 2009). Lecithotrophy could have evolved through the accumulation of small-effect changes throughout the genome, but our results suggest that only one or a few genes (genomic regions) of large effect can account for major reduction in the length of larval chaetae. A complete accounting of this trait’s evolution requires genetic dissection of the maternal effects, which awaits analysis of F_3_ and subsequent generations.

Reconciling the phenoptypic results from F_1_ and F_2_ populations with the persistence of poecilogony is challenging. Our results show that random mating in *S. benedicti* would result in heterogeneous populations that include intermediate larval phenotypes, which are absent from most natural populations (with rare exceptions: Levin and Huggett, 1990). Our study and others (Levin *et al*., 1991; Schulze *et al*., 2000) show that planktotrophic and lecithotrophic worms can freely mate in the lab. Further, gene flow is occurring in natural populations (Zakas and Wares, 2012). The ultimate explanation may involve ecologically structured metapopulation dynamics and directional gene flow (Zakas and Hall, 2012), and discovery of the causal genes for lecithotrophy will facilitate our dissection of its ecological genetic basis. In the meantime, our data show that a fraction of F_1_ larvae are indistinguishable in gut length and chaetal traits from their pure-morph parents, with the variable penetrance of chaetal growth playing a central role. In LxB crosses (lecithotrophic mothers and planktotrophic fathers), ∼30% of F_1LB_ larvae are phenotypically similar to LL larvae. Assuming these individuals are subject to the same selection pressures as LL larvae, this could account for gene flow in the face of strong selection against intermediate phenotypes. F_1BL_ larvae (planktotrophic mothers and lecithotrophic fathers) have less phenotypic overlap with BB larvae, but some overlap occurs. Occasional F_1_ larval migrants could provide the gene flow required to homogenize the genomic background among populations while strong selection preserves differentiation at the loci directly responsible for larval phenotypes. Genetic analysis offers the prospect of eventually discovering those loci.

## Acknowledgements

We thank Bruno Pernet for sending us animals he collected in Long Beach, Jenn Deutscher for assistance with animal husbandry and imaging, Max Bernstein, Vicky Cattani, Annalise Paaby, Taniya Kaur and Luke Noble for comments on the manuscript, and NYU, the Zegar Family Foundation and NSF grant IOS-1350926 for financial support.

